# Transcriptomic and proteomic profiling of Na_V_1.8-expressing mouse nociceptors

**DOI:** 10.1101/2022.07.14.499815

**Authors:** Manuela Schmidt, Julia Regina Sondermann, David Gomez-Varela, Queensta Millet, John N Wood, Jing Zhao

**Author notes:** **Correspondence:** Manuela Schmidt; John N Wood and Jing Zhao.

## Abstract

Nociceptors play an essential role in both acute pain and chronic pain conditions. and have recently been classified into distinct subsets using single-cell transcriptional profiling. In this study, we examined protein levels in dorsal root ganglia using DIA Mass-spectrometry technologies with Na_V_1.8Cre^+/-^; ROSA26-flox-stop-flox-DTA (Diphtheria toxin fragment A) mutant mice (Na_V_1.8Cre-DTA), in which Na_V_1.8-expressing neurons (mainly nociceptors) in dorsal root ganglia (DRG) were ablated. The results show that 353 transcripts and 78 proteins, including nociceptor-specific sodium channels Na_V_1.8 (Scn10a) and NaV1.9 (Scn11a), were specifically expressed in nociceptors of DRG. A comparative analysis revealed that about 40% of nociceptor-specific proteins are shared within the nociceptor-specific transcript dataset. Scatter plots show that the proteome and transcriptome datasets in nociceptors have a moderate correlation (*r* = 0.4825), indicating the existence of post-transcriptional and post-translational gene regulation in nociceptors. This combined profiling study provides a unique resource for sensory studies, especially for pain research.

## INTRODUCTION

Chronic pain is poorly treated in the clinic as most available analgesics have low efficacy and can cause serious side effects. A recent survey on chronic pain shows that 1 in 5 European people suffers from chronic pain, thus, the development of new types of analgesic drugs is urgently needed (Alliance, 2017). This depends on a more detailed understanding of the molecular mechanisms underlying pain syndromes. The concept of nociceptors was introduced by neurophysiologist Sir Charles Sherrington in the early 20th century and their existence demonstrated by Edward R. Perl (Mason, 2007; Sherrington, 1903; Wood, 2020). Specialized primary sensory neurons resident in dorsal root ganglia (DRG) and trigeminal ganglia play a fundamental role in both acute pain and chronic pain conditions (Abrahamsen et al., 2008; Dubin & Patapoutian, 2010; Middleton et al., 2021; Reichling & Levine, 2009). DRG are located outside the blood-brain barrier rendering them an important therapeutic target of chronic pain, in particular for peripherally-acting treatments (Price et al., 2018; Sapunar et al., 2012; Woolf & Ma, 2007). Expression of the sodium channel Na_V_1.8 defines a set of nociceptors involved in mechanical and inflammatory pain (Akopian et al., 1999; Wood, 2020).

In past decades, many studies have focused on investigating pain-associated gene expression changes in DRG. Microarray profiling and RNA-Seq have been widely employed to investigate the DRG transcriptome in many species such as mouse, rat, primate, and human (Barry et al., 2018; Chiu et al., 2014; Gong et al., 2016; J. Kupari et al., 2021; Li et al., 2016; Megat, Ray, Moy, et al., 2019; Megat, Ray, Tavares-Ferreira, et al., 2019; Ray et al., 2018; Yokoyama et al., 2020). Among these reports, some assessed mRNA levels specifically in Na_V_1.8-expressing (Na_V_1.8^+^) nociceptors.

For example, Thakur *et al*. identified 920 transcripts enriched in nociceptors using Na_V_1.8-tdTomato mice combined with magnetic cell sorting (MACS) and RNA-Seq technologies (Thakur et al., 2014). In 2008, we described insights into the nociceptor transcriptome using microarrays with Na_V_1.8Cre^+/-^; ROSA26-flox-stop-flox-DTA (Diphtheria toxin fragment A) mutant mice (DTA) (Abrahamsen et al., 2008), in which Na_V_1.8^+^-expressing DRG neurons, representing mainly nociceptors, were specifically ablated by expression of DTA. More recently, subtypes of DRG neurons, including nociceptors, were identified, and distinguished by comprehensive single-cell RNA-Seq profiles in mouse and primates generating a highly valuable reference atlas of gene expression in DRG and beyond (Jussi Kupari et al., 2021; Usoskin et al., 2015; Zeisel et al., 2018). Yet, transcript levels only show limited correspondence with protein abundance given diverse cellular buffering mechanisms, e.g. regulation at the level of translation and post-translationally (Liu et al., 2016; Reimegård et al., 2021). Furthermore, Megat *et al*. reported the translatome of Na_V_1.8^+^ DRG neurons using translating ribosome affinity purification (TRAP) and RNA-Seq technologies (Megat, Ray, Moy, et al., 2019). Importantly, they observed only minor correlations between the translatome and transcriptome. In functional terms, however, proteins are the building blocks of a cell, and they are crucially implicated in determining phenotypes, including pain-related outcomes and behaviours. Nonetheless, compared to the transcriptome of nociceptors, the Na_V_1.8^+^ nociceptor proteome remains undefined.

We have recently shown the immense potential of defining pain-associated proteome dynamics in DRG by data-independent acquisition mass spectrometry (DIA-MS) (Barry et al., 2018; De Clauser et al., 2022; Rouwette et al., 2016). Here, we used DIA-MS to reveal the previously unknown protein set-up of nociceptors by using aforementioned DTA mice (Abrahamsen et al., 2008). Following the workflow chart shown in Figure 1, we, in parallel, re-analysed the raw microarray data (ArrayExpress: E-MEXP-1622) obtained in our previous study (Abrahamsen et al., 2008) with the latest version of Transcriptome Analysis Console (TAC) Software 4.0.2. Furthermore, the two datasets were compared and are presented here with volcano plots, Venn diagrams, Pearson scatter plots and bar charts. The top 50 nociceptor-enriched transcripts/proteins can be found in Supplementary Tables 1 and 2. Overall, this study provides a valuable resource atlas covering transcripts and proteins in Na_V_1.8^+^ nociceptors of mouse DRG.

**Figure 1.**
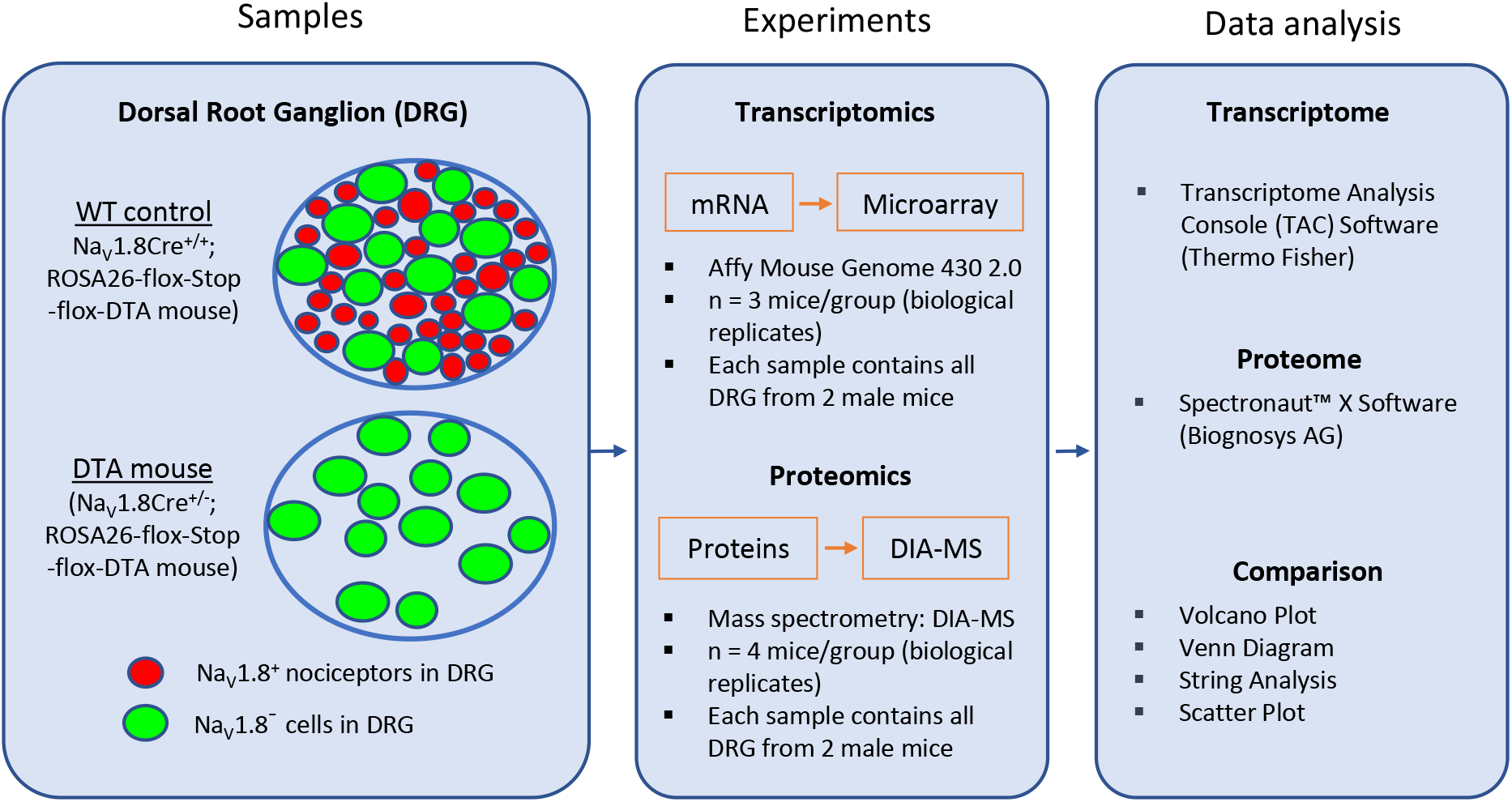
Schematic diagram for transcriptome and proteome analysis in Na_V_1.8-expressing (Na_V_1.8^+^) DRG neurons (mainly nociceptors). The diagram shows the key steps of the workflow. First, RNA and protein samples from DRG were collected from both Na_V_1.8Cre^+/-^; ROSA26-flox-Stop-flox-DTA mutant mice (DTA), in which Na_V_1.8^+^ nociceptors have been ablated, and Na_V_1.8Cre^+/+^; ROSA26-flox-stop-flox-DTA littermates as “wild-type” control mice (WT). Isolated RNA and proteins were submitted to Affymetrics microarrays and data independent acquisition-mass spectrometry (DIA-MS), respectively. Generated datasets were analysed using Transcriptome Analysis Console (TAC) software for transcriptome profiling, and Spectronaut™ Software (Biognosys AG) for proteome profiling.

## MATERALS AND METHODS

### Animals

To ablate the neuronal population of DRG neurons expressing Na_V_1.8, heterozygous Na_V_1.8Cre^+/-^ mice (Nassar et al., 2004) were crossed to homozygous Cre-dependent Diphtheria Toxin fragment A (ROSA26-flox-stop-flox-DTA) mice (Abrahamsen et al., 2008; Ivanova et al., 2005). In the offspring, Na_V_1.8Cre^+/-^; ROSA26-flox-stop-flox-DTA mice (DTA) and Na_V_1.8Cre^+/+^; ROSA26-flox-stop-flox-DTA littermates wild-type mice (WT) were used as nociceptor-deficient mice and “wild-type” control mice, respectively (Figure 1). The genotyping analysis was performed as previously descripted (Abrahamsen et al., 2008).

### mRNA Microarray

The mRNA Microarray experiment was performed as described in our previous study (Abrahamsen et al., 2008). Raw data were stored in ArrayExpress (E-MEXP-1622) and Transcriptome Analysis Console (TAC) Software (version 4.0.2) was used for extracting the raw data. A list of all identified transcripts can be found in Supplementary Table 7.

### Sample preparation for DIA-MS

DRG were isolated from eight DTA males, aged between 8-12-weeks-old, and eight littermate WT male controls, i.e., four biological replicates with two mice/replicate in total. Protein samples of DRG were prepared as described previously (Barry et al., 2018). All reagents were obtained from Roth. All DRG samples were stored at -80 degrees until further use.

### DIA-MS and data analysis

All solvents were HPLC-grade from Sigma-Aldrich and all chemicals were obtained from Sigma-Aldrich if not stated otherwise. All steps of DIA-MS and its analysis were performed by Biognosys AG (Zuerich, Switzerland) essentially as described (Bruderer et al., 2015; Rouwette et al., 2016) with the following modifications: For Global HRM profiling, 2 µg of peptides were injected via an in-house packed C18 column (Dr. Maisch ReproSil Pur, 1.9 µm particle size, 120 Åpore size; 75 µm inner diameter, 50 cm length, New Objective) on a Thermo Scientific Easy nLC 1200 nano-liquid chromatography system connected to a Thermo Scientific Fusion Lumos Tribrid mass spectrometer equipped with a standard nano-electrospray source. LC solvents were A: 1 % acetonitrile in water with 0.1 % FA; B: 15 % water in acetonitrile with 0.1 % FA. The nonlinear LC gradient was 1 - 55 % solvent B in 120 minutes followed by 55 – 90 % B in 10 seconds, 90 % B for 10 minutes, 90 % - 1 % B in 10 seconds and 1 % B for 5 minutes. A DIA method with one full range survey scan and 40 DIA windows was adopted from a previous study (Bruderer et al., 2015). HRM mass spectrometric data were analyzed using Spectronaut software (Biognosys). The false discovery rate on peptide and protein level was set to 1 %, data was filtered using row-based extraction. Data quality was analyzed using directDIA analysis using Spectronaut X and the normalization applied in Spectronaut. Data analysis was performed using mouse UniProt fasta downloaded 2018-07-01. We used two analysis pipelines as described previously (De Clauser et al., 2022): the spectral library-based search and the directDIATM workflow developed at Biognosys using Biognosys Factory settings in Spectronaut X. Label-free Quantitation was executed on MS2-level using the area under the curve and data were filtered by Q-value sparse (precursors robustly found in at least one sample). Statistical testing of differential protein abundances between conditions was calculated in Spectronaut for each protein ID by performing a pairwise *t*-test. Benjamini-Hochberg (BH)-adjusted P-values (Q-values) were used for multiple testing and significantly altered proteins were defined by setting a cut-off of Q < 0.05. Significantly regulated proteins from both analysis pipelines were pooled in a combined candidate list and duplicates as well as single-peptide-hits were removed. Any potential keratin and serum albumin contaminations were also removed. A list of all quantified proteins can be found in Supplementary Table 8.

All raw data and sample report files have been deposited to the ProteomeXchange Consortium via the PRIDE partner repository (Perez-Riverol *et al*., 2019) with the dataset identifier PXD034447.

### Data analysis and statistics

Raw data from mRNAs Microarray and protein DIA-MS were analysed with TAC software (4.0.2) and Spectronaut X software, respectively. Further data analysis was performed with GraphPad Prism 9 to generate volcano plots, Venn diagrams, bar graphs, and scatter plots for comparative data analysis. All symbols of transcripts and proteins were converted to GMI approved gene symbols for ease of comparison among the two datasets.

All values are presented as Mean ± S.E.M. Data were analysed by two-way ANOVA and Student’s *t-*test. Differences were considered significant at FDR adjusted P < 0.05 for transcriptome data. For proteome analysis, Benjamini-Hochberg (BH)-adjusted P-values were used for multiple testing and significantly altered proteins were defined by setting a cut-off of Q < 0.05 as previously described (De Clauser et al., 2022).

### Transcriptomic profiling of DRG

To study the gene expression at mRNA level in DRG, especially in Na_V_1.8^+^ neurons (mainly nociceptors) (Stirling et al., 2005), we re-analysed the raw DNA microarray data (Abrahamsen et al., 2008) using the latest version (4.0.2) of Transcriptome Analysis Console (TAC) software (Thermo Fisher Scientific). If one transcript was represented by multiple entries, these were removed to obtain a list, in which each transcript is represented by one entry. Finally, 21,037 transcripts were identified in DRG, among which 353 (1.68% of total transcripts) were significantly down-regulated (Fold-Change, FC < -2.0; FDR adjusted P*-*value < 0.05) in DTA mice (Figure 2A), suggesting that these transcripts were enriched in Na_V_1.8^+^ nociceptors in DRG. As expected, both nociceptive sodium channels Na_V_1.8 (*Scn10a*) and NaV1.9 (*Scn11a*) were highly enriched in nociceptors (Figure 2A). In contrast, 207 transcripts (0.98% of total transcripts) were up-regulated (FC > 2.0; FDR adjusted P*-*value < 0.05) in DRG in DTA mice, indicating that those transcripts exhibit very low to no expression in Na_V_1.8^+^ neurons. Rather, these are primarily expressed in Na_V_1.8^-^ cells including large-diameter DRG neurons and/or satellite glial cells of DRG. The top 50 transcripts, which are enriched in Na_V_1.8^+^ DRG neurons are listed in Supplementary Table 1. A list of all identified transcripts can be found in Supplementary Table 7.

**Figure 2.**
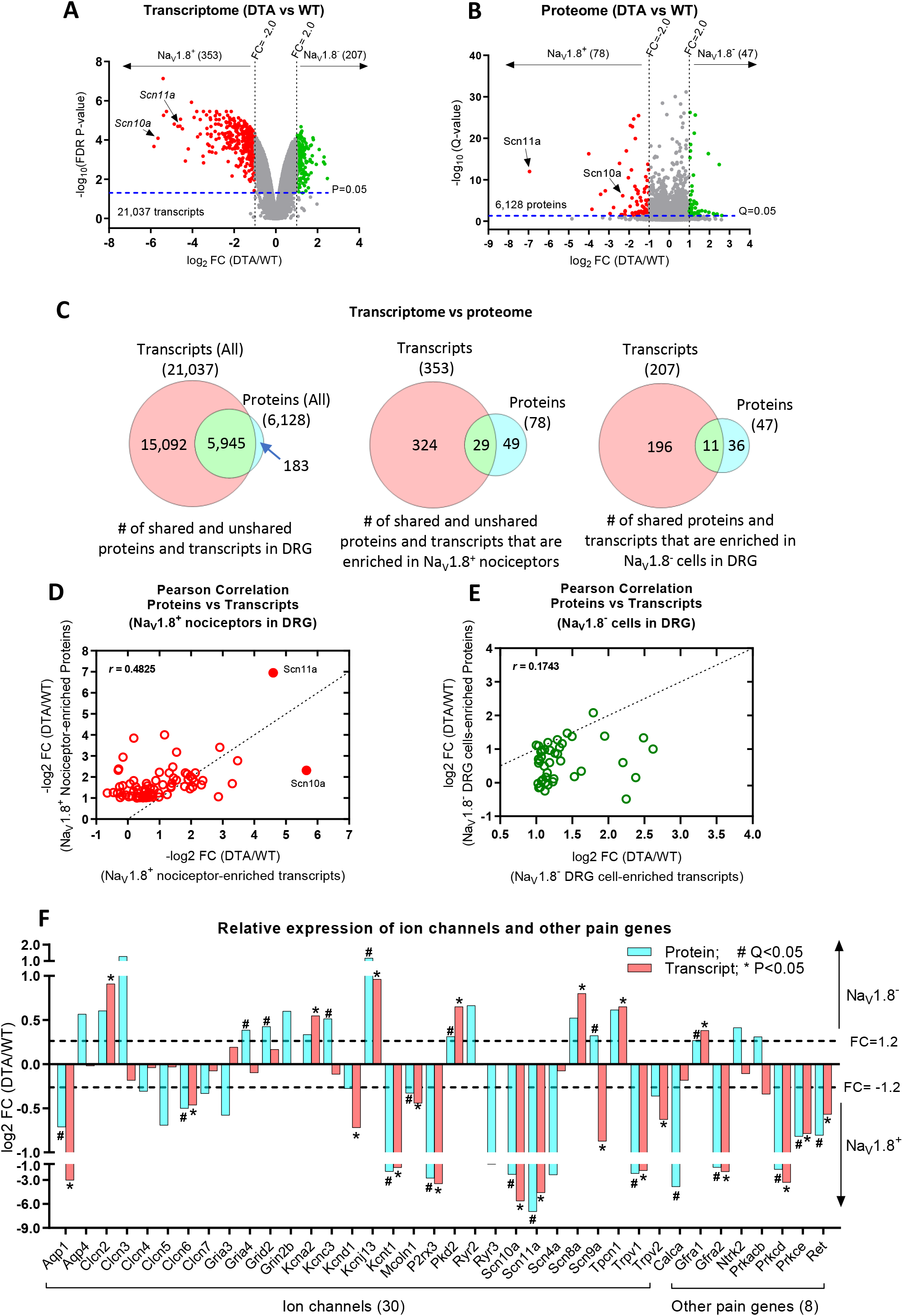
Profiling and comparison of the transcriptome and proteome of DRG from DTA mice and littermate control mice. **(A)** Volcano plot shows the average fold-change (log_2_FC) in expression levels of transcripts versus P-values (log_10_FDR P-value). Results are filtered by FC: FC < -2.0, red dots; FC > 2.0, green dots; all with P*-*value < 0.05. **(B)** Volcano plot shows the average log_2_FC in relative protein abundance versus Q-values (log_10_; Q-values represent P-values corrected for multiple-testing, please see methods for details). Results are filtered by FC: FC < -2.0, red dots; FC > 2.0, green dots; all with Q*-*value < 0.05. **(C)** Venn diagrams indicate the numbers of transcripts and proteins quantified in DRG (left panel), enriched in Na_V_1.8^+^ DRG neurons, mainly nociceptors (middle panel), and enriched in Na_V_1.8^-^ DRG cells (right panel), non-nociceptors. **(D-E)** Scatter Plots (Pearson correlation) show the correlation between proteome and transcriptome data: moderate correlation (*r* = 0.4825) in Na_V_1.8^+^ nociceptors (D) and low correlation (*r* = 0.1743) in Na_V_1.8^-^ cells of DRG (E). **(F)** Bar graphs show the comparison of protein versus transcript levels (log_2_FC) of a selection of ion channels and other pain genes either enriched in Na_V_1.8^+^ nociceptors or in Na_V_1.8^-^ DRG cells (DTA versus WT). * P-value < 0.05 in the transcriptome dataset; # Q-value < 0.05 in the proteome dataset.

### Proteomic profiling of DRG

Next, we deeply profiled the proteome of DRG neurons using our established DIA-MS workflow as previously described (Barry et al., 2018). In total, we quantified 6,128 proteins among which 78 (0.13% of total proteins) were significantly down-regulated (FC < -2.0; Q*-*value < 0.05) in DTA mice (Figure 2B), indicating that these 78 proteins (Supplementary Table 4) were enriched in Na_V_1.8^+^ DRG neurons. In line with our transcriptome analysis, Na_V_1.8 (*Scn10a*) and NaV1.9 (*Scn11a*) appeared to be highly enriched in nociceptors. In contrast, 47 proteins (Supplementary Table 5) (0.77% of total proteins) were significantly up-regulated (FC > 2.0; *Q-*value < 0.05) in DRG of DTA mice, suggesting that these proteins are primarily expressed in Na_V_1.8^-^ cells in DRG (likely large DRG neurons and/or satellite cells). The top 50 proteins, which are enriched in Na_V_1.8^+^ DRG neurons, are listed in Supplementary Table 2. A list of all quantified proteins can be found in Supplementary Table 7.

### Comparison of transcriptome and proteome of the nociceptors

We then compared the proteome and transcriptome datasets as shown in Venn diagrams (Figure 2C) and Pearson correlation scatter plots (Figure 2D and 2E). The left Venn diagram (left panel in Figure 2C) shows that 97% (5,945 out of 6,128) proteins could be matched to their corresponding transcripts. The remaining unmatched 183 proteins are listed in Supplementary Table 3. As expected, the transcriptome dataset is much more comprehensive than the protein dataset owing to technical reasons. Consequently, we did not obtain protein abundance data on 2/3 of detected transcripts, including pain-relevant genes such as TrpA1 and TrkA. The middle panel in Figure 2C demonstrates that only 29 out of 78 (37.2%) proteins enriched in Na_V_1.8^+^ DRG neurons appeared to be nociceptor-enriched on the mRNA-level. Similarly, the overlap of non-nociceptor-enriched proteins and transcripts was relatively low: only 11 out of 47 (23.4%) proteins could be matched on the transcript level (Supplementary Table 4 and 5). To directly assess the correlation between our transcript and protein datasets, we calculated Pearson’s correlation coefficients with GraphPad Prism 9 software. The results show that the two datasets have moderate correlation (*r* = 0.4825) in Na_V_1.8^+^ neurons (Figure 2D) and low correlation (*r* = 0.1743) in non-nociceptor Na_V_1.8^-^ cells in DRG (Figure 2E). In a next step, we compared transcript and protein levels of a selection of ion channels and known pain genes. For the most part transcript and protein data were well correlated in respect to their enrichment in Na_V_1.8^+^ nociceptors and Na_V_1.8^-^ non-nociceptors, respectively. For example, Na_V_1.8 (*Scn10a*), NaV1.9 (*Scn11a*), *Trpv1, P2rx3*, PKC Delta (*Prkcd*) and PKC Epsilon (*Prkce*), c-RET (*RET*), GDNF Receptor A2 (*Gfra2*) and Aquaporin 1 (*Aqp1*) were significantly enriched in Na_V_1.8^+^ nociceptors (Figure 2F), while potassium channels Kcna2 and Kcnj13, Polycystin-2 (Pkd2), Scn8a, and GDNF Receptor A1 (*Gfra1*) were enriched in Na_V_1.8^-^ cells of DRG. In contrast, some molecules showed opposite trends of transcript and protein levels, such as NaV1.7 (*Scn9a*) and PKA Beta (*Prkacb*), a finding that awaits further investigation by other methods. To validate our findings we assessed the expression of 5 selected transcripts/proteins (Scn10a, Scn11a, P2rx3, Prkcd and prostate acid phosphatase, Accp), which are shared among Top-50 transcripts (Supplementary Table 1) and Top-50 proteins (Supplementary Table 2), by comparison to a single cell RNA-Seq database (www.mousebrain.org) (Zeisel et al., 2018). This comparison confirmed here shown enrichment in nociceptors (Supplementary Table 6).

## CONCLUSION

This comparative profiling study on gene expression at both mRNA and protein levels in Na_V_1.8^+^ nociceptors in mice provides a unique and highly valuable source for future studies on the molecular basis of somatosensation and pain.

## Supporting information

Supplementary Table 7 - Transcriptome

Supplementary Table 8 - Proteome

## DATA AVAILABILITY STATEMENT

Transcriptome data are publicly accessible at ArrayExpress linked at https://www.ebi.ac.uk/arrayexpress/experiments/E-MEXP-1622. All proteome raw data and sample report files have been deposited to the ProteomeXchange Consortium via the PRIDE partner repository (Perez-Riverol et al., 2019) with the dataset identifier PXD034447.

## ETHICS STATEMENT

All animal procedures were approved by University College London Research Ethics Committee and conformed to UK Home Office regulations.

## AUTHOR CONTRIBUTIONS

MS, JW and JZ were involved in conceptualization, formal analysis, investigation, methodology and writing the manuscript. DGV, JRS, and JZ were involved in investigation and data collection. MQ was involved in investigation. MS, JW and JZ were involved in data analysis. All authors contributed to data discussion and read approved the manuscript.

## CONFLICT OF INTEREST

The authors declare that the research was conducted in the absence of any commercial or financial relationships that could be construed as a potential conflict of interest.

### ACKNOWLEDGMENTS

We thank Mr. Sam Gossage for handling the samples, and Prof James J Cox for his valuable suggestions on the project and manuscript. We thank Biognosys AG for performing mass spectrometry.

## SUPPLEMENTARY MATERIAL

Supplementary Materials for this article can be found in separate files. A list of these files shows below.

Supplementary Table 1. Top-50 transcripts enriched in Na_V_1.8^+^ nociceptors of DRG

Supplementary Table 2. Top-50 proteins enriched in Na_V_1.8^+^ nociceptors of DRG

Supplementary Table 3. Unmatched proteins (183) for which no corresponding transcripts could be found in the transcript list

Supplementary Table 4. Na_V_1.8^+^ nociceptor-enriched proteins (78) in DRG

Supplementary Table 5. Proteins (47) enriched in Na_V_1.8^-^ cells of DRG

Supplementary Table 6. The expression (as indicated by MouseBrain.org) of 5 selected candidates shared among Top-50 transcripts (Suppl. Table 1) and Top-50 proteins (Suppl. Table 2)

Supplementary Table 7. All transcripts detected in mouse DRG using mRNA Microarray

Supplementary Table 8. All proteins detected in mouse DRG using DIA-MS

