## Supplementary Table 1-6 for "Transcriptomic and proteomic profiling of Na_V_1.8-expressing mouse nociceptors"

### SUPPLEMENTARY MATERIAL

Supplementary Table 1. Top-50 transcripts enriched in $Na_V1.8^+$ nociceptors

Supplementary Table 2. Top-50 proteins enriched in $Na_V1.8^+$ nociceptors

Supplementary Table 3. Unmatched proteins (183) for which no corresponding transcripts could be found in the transcript list

Supplementary Table 4. $Na_V1.8^+$ nociceptor-enriched proteins (78) in DRG

Supplementary Table 5. Proteins (47) -enriched in $Na_V1.8^-$ cells of DRG

Supplementary Table 6. The expression (as indicated by MouseBrain.org) of 5 representative candidates shared among Top-50 transcripts (Suppl. Table 1) and Top-50 proteins (Suppl. Table 2)

Supplementary Table 7. All transcripts (21,037) detected in mouse DRG using mRNA Microarray

Supplementary Table 8. All proteins (6,128) detected in mouse DRG using DIA-MS

### Suppl. Table 1 | Top 50 transcripts enriched in $Na_V1.8^+$ nociceptors

| No. | MGI Symbol | Description | Probe ID | FDR P-val | FC (DTA/WT) |
|---|---|---|---|---|---|
| 1 | Tmem45b | transmembrane protein 45b | 1424357_at | 2.15E-04 | -57.3 |
| 2 | Scn10a | sodium channel, voltage-gated, type X, alpha | 1450266_at | 8.16E-05 | -50.1 |
| 3 | Lpar3 | lysophosphatidic acid receptor 3 | 1418723_at | 7.43E-08 | -42.4 |
| 4 | Mal2 | mal, T cell differentiation protein 2 | 1427042_at | 5.58E-06 | -41.6 |
| 5 | Gna14 | guanine nucleotide binding protein, alpha 14 | 1420385_at | 3.53E-06 | -37.8 |
| 6 | Kcnmb1 | potassium large conductance calcium-activated channel, subfamily M, beta member 1 | 1421400_at | 1.54E-05 | -29.2 |
| 7 | Nppb | natriuretic peptide type B | 1450791_at | 2.01E-05 | -26.0 |
| 8 | Scn11a | sodium channel, voltage-gated, type XI, alpha | 1420784_at | 1.97E-05 | -24.2 |
| 9 | Ly86 | lymphocyte antigen 86 | 1422903_at | 8.73E-06 | -23.8 |
| 10 | Npy2r | neuropeptide Y receptor Y2 | 1417489_at | 2.72E-05 | -22.3 |
| 11 | Trpa1 | transient receptor potential cation channel, subfamily A, member 1 | 1457164_at | 1.18E-03 | -20.2 |
| 12 | Mrgpra3 | MAS-related GPR, member A3 | 1426121_at | 2.86E-04 | -18.4 |
| 13 | Grik1 | glutamate receptor, ionotropic, kainate 1 | 1427676_a_at | 1.19E-06 | -16.5 |
| 14 | Gm16364 | predicted gene 16364 | 1441647_at | 7.11E-06 | -15.1 |
| 15 | Celf6 | CUGBP, Elav-like family member 6 | 1429790_at | 3.53E-06 | -13.8 |
| 16 | Rasgrp1 | RAS guanyl releasing protein 1 | 1421176_at | 6.98E-05 | -13.4 |
| 17 | Ctxn3 | cortexin 3 | 1438059_at | 9.01E-06 | -12.3 |
| 18 | Mrgpra2a | MAS-related GPR, member A2A; MAS-related GPR, member A2B | 1451858_at | 5.03E-05 | -12.3 |
| 19 | Slc16a12 | solute carrier family 16 (monocarboxylic acid transporters), member 12 | 1434188_at | 6.73E-04 | -11.2 |
| 20 | P2rx3 | purinergic receptor P2X, ligand-gated ion channel, 3 | 1425093_at | 3.53E-06 | -11.1 |
| 21 | Th | tyrosine hydroxylase | 1420546_at | 1.48E-03 | -11.0 |
| 22 | Syt9 | synaptotagmin IX | 1423258_at | 5.26E-06 | -10.0 |
| 23 | Prkcd | protein kinase C, delta | 1422847_a_at | 2.57E-05 | -9.9 |
| 24 | Otoa | otoancorin | 1428037_at | 4.90E-05 | -9.7 |
| 25 | Ceacam10 | carcinoembryonic antigen-related cell adhesion molecule 10 | 1417074_at | 1.62E-04 | -9.6 |
| 26 | Mab21l1 | mab-21-like 1 (C. elegans) | 1421369_a_at | 1.92E-04 | -9.1 |
| 27 | Rgs8 | regulator of G-protein signaling 8 | 1453060_at | 3.91E-04 | -9.1 |
| 28 | Rab27b | RAB27B, member RAS oncogene family | 1417214_at | 3.53E-06 | -9.0 |
| 29 | Cd44 | CD44 antigen | 1423760_at | 7.49E-06 | -8.9 |
| 30 | Trpc6 | transient receptor potential cation channel, subfamily C, member 6 | 1449431_at | 9.01E-06 | -8.7 |
| 31 | Nrn1l | neuritin 1-like | 1456819_at | 6.62E-05 | -8.6 |
| 32 | Il31ra | interleukin 31 receptor A | 1451535_at | 2.07E-05 | -8.6 |
| 33 | S100a7l2 | S100 calcium binding protein A7 like 2 | 1445608_at | 5.18E-05 | -8.6 |
| 34 | Ntrk1 | neurotrophic tyrosine kinase, receptor, type 1 | 1443990_at | 4.16E-04 | -8.3 |
| 35 | Aqp1 | aquaporin 1 | 1416203_at | 2.31E-04 | -8.2 |
| 36 | Acpp | acid phosphatase, prostate | 1419832_s_at | 3.68E-05 | -7.5 |
| 37 | Tmem158 | transmembrane protein 158 | 1428074_at | 5.85E-06 | -7.5 |
| 38 | Paqr5 | progestin and adipoQ receptor family member V | 1456152_at | 1.54E-05 | -7.4 |
| 39 | Ms4a3 | membrane-spanning 4-domains, subfamily A, member 3 | 1420572_at | 4.58E-04 | -7.4 |
| 40 | Klk5 | kallikrein related-peptidase 5 | 1429230_at | 9.01E-06 | -7.3 |
| 41 | Prkcq | protein kinase C, theta | 1426044_a_at | 9.27E-05 | -7.3 |
| 42 | Synpr | synaptoporin | 1423640_at | 3.53E-06 | -7.2 |
| 43 | Ccdc68 | coiled-coil domain containing 68 | 1439109_at | 4.15E-06 | -7.1 |
| 44 | Gpr35 | G protein-coupled receptor 35 | 1449976_a_at | 1.56E-05 | -6.7 |
| 45 | Far2 | fatty acyl CoA reductase 2 | 1435621_at | 5.03E-05 | -6.6 |
| 46 | Capn9 | calpain 9 | 1422876_at | 8.07E-05 | -6.6 |
| 47 | Trpc3 | transient receptor potential cation channel, subfamily C, member 3 | 1417577_at | 9.25E-06 | -6.6 |
| 48 | Agtr1a | angiotensin II receptor, type 1a | 1436739_at | 5.11E-06 | -6.5 |
| 49 | Fam184b | family with sequence similarity 184, member B | 1421595_at | 1.77E-05 | -6.2 |
| 50 | Nmb | neuromedin B | 1419405_at | 4.54E-05 | -5.9 |

MGI gene symbols, gene names, Affymetrix probe IDs, FDR adjusted P-values and fold changes (FC) are shown. FC is the ratio (DTA/WT) of the averaged relative transcript abundance in DRG. (-) indicates enrichment in $Nav1.8^+$ nociceptors. Transcripts were sorted by FC.

### Suppl. Table 2 | Top 50 proteins enriched in $Na_V1.8^+$ nociceptors

| No. | MGI Symbol | Description | UniProt IDs | Q-value | FC (DTA/WT) |
|---|---|---|---|---|---|
| 1 | Scn11a | sodium channel, voltage-gated, type XI, alpha | Q9R053 | 1.01E-12 | -124.0 |
| 2 | Camk2a | calcium/calmodulin-dependent protein kinase II alpha | F8WIS9 | 5.47E-17 | -16.0 |
| 3 | Calca | calcitonin/calcitonin-related polypeptide, alpha | Q99JA0 | 1.36E-03 | -14.4 |
| 4 | Acpp | acid phosphatase, prostate | Q8CE08 | 3.20E-07 | -10.7 |
| 5 | Rgs3 | regulator of G-protein signaling 3 | Q9DC04 | 4.55E-08 | -9.1 |
| 6 | Kndc1 | kinase non-catalytic C-lobe domain (KIND) containing 1 | Q0KK55 | 1.49E-02 | -7.7 |
| 7 | P2rx3 | purinergic receptor P2X, ligand-gated ion channel, 3 | A2AW03 | 4.79E-04 | -6.8 |
| 8 | Dgki | diacylglycerol kinase, iota | D3YWQ0 | 1.17E-14 | -5.6 |
| 9 | Tmem177 | transmembrane protein 177 | Q8BPE4 | 2.59E-02 | -5.2 |
| 10 | Chsy3 | chondroitin sulfate synthase 3 | Q5DTK1 | 2.72E-02 | -5.0 |
| 11 | Scn10a | sodium channel, voltage-gated, type X, alpha | Q6QIY3 | 7.17E-07 | -5.0 |
| 12 | Slc36a1 | solute carrier family 36 (proton/amino acid symporter), member 1 | Q8K4D3 | 3.57E-02 | -4.7 |
| 13 | Arhgap28 | Rho GTPase activating protein 28 | E9Q642 | 4.72E-03 | -4.6 |
| 14 | Ptrh1 | peptidyl-tRNA hydrolase 1 homolog | Q8BW00 | 2.95E-02 | -4.6 |
| 15 | Trpv1 | transient receptor potential cation channel, subfamily V, member 1 | Q704Y3 | 3.12E-03 | -4.6 |
| 16 | Phf24 | PHD finger protein 24 | Q80TL4 | 1.13E-17 | -4.4 |
| 17 | Nedd4l | neural precursor cell expressed, developmentally down-regulated gene 4-like | E9PXB7 | 3.41E-11 | -4.1 |
| 18 | Kcnt1 | potassium channel, subfamily T, member 1 | A0A0G2JER3 | 2.66E-04 | -3.9 |
| 19 | Dgkh | diacylglycerol kinase, eta | A0A2I3BQ43 | 8.18E-24 | -3.8 |
| 20 | Dgkz | diacylglycerol kinase zeta | A2AHJ7 | 4.34E-13 | -3.7 |
| 21 | Pirt | phosphoinositide-interacting regulator of transient receptor potential channels | Q8BFY0 | 2.51E-06 | -3.6 |
| 22 | Disp2 | dispatched RND tramsporter family member 2 | Q8CIP5 | 1.06E-02 | -3.6 |
| 23 | Eml1 | echinoderm microtubule associated protein like 1 | Q05BC3 | 1.66E-23 | -3.6 |
| 24 | AU040320 | expressed sequence AU040320 | Q8K135 | 4.55E-02 | -3.6 |
| 25 | Osbpl3 | oxysterol binding protein-like 3 | D3YTT6 | 2.25E-25 | -3.4 |
| 26 | Scg3 | secretogranin III | P47867 | 3.86E-09 | -3.3 |
| 27 | D130043K22Rik | RIKEN cDNA D130043K22 gene | Q5SZV5 | 8.33E-03 | -3.2 |
| 28 | Prkcd | protein kinase C, delta | P28867 | 1.22E-20 | -3.2 |
| 29 | Trarg1 | trafficking regulator of GLUT4 (SLC2A4) 1 | Q8C838 | 4.56E-02 | -3.2 |
| 30 | Clgn | calmegin | P52194 | 2.23E-08 | -3.1 |
| 31 | Dgkg | diacylglycerol kinase, gamma | Q91WG7 | 1.78E-05 | -3.1 |
| 32 | Galnt17 | polypeptide N-acetylgalactosaminyltransferase 17 | Q7TT15 | 9.92E-05 | -3.0 |
| 33 | St8sia3 | ST8 alpha-N-acetyl-neuraminide alpha-2,8-sialyltransferase 3 | Q64689 | 6.97E-03 | -3.0 |
| 34 | Fxyd2 | FXYD domain-containing ion transport regulator 2 | Q04646 | 2.12E-07 | -2.9 |
| 35 | Rgs10 | regulator of G-protein signalling 10 | Q9CQE5 | 5.52E-08 | -2.9 |
| 36 | Srl | sarcalumenin | Q7TQ48 | 2.91E-02 | -2.9 |
| 37 | Dolpp1 | dolichyl pyrophosphate phosphatase 1 | Q9JMF7 | 3.47E-02 | -2.9 |
| 38 | Plcb3 | phospholipase C, beta 3 | P51432 | 3.43E-26 | -2.9 |
| 39 | Plcxd3 | phosphatidylinositol-specific phospholipase C, X domain containing 3 | G3X9A7 | 4.50E-03 | -2.8 |
| 40 | Atrnl1 | attractin like 1 | Q6A051 | 6.51E-03 | -2.8 |
| 41 | Gfra2 | glial cell line derived neurotrophic factor family receptor alpha 2 | O08842 | 1.94E-06 | -2.7 |
| 42 | Kctd16 | potassium channel tetramerisation domain containing 16 | Q5DTY9 | 4.53E-03 | -2.7 |
| 43 | Ids | iduronate 2-sulfatase | Q08890 | 1.21E-05 | -2.6 |
| 44 | Myh4 | myosin, heavy polypeptide 4, skeletal muscle | Q5SX39 | 3.93E-03 | -2.6 |
| 45 | Ccdc141 | coiled-coil domain containing 141 | A2AST1 | 2.43E-04 | -2.5 |
| 46 | Cpt1c | carnitine palmitoyltransferase 1c | Q8BGD5 | 1.35E-02 | -2.5 |
| 47 | Hps1 | HPS1, biogenesis of lysosomal organelles complex 3 subunit 1 | A0A0R4J062 | 3.12E-02 | -2.4 |
| 48 | Myh2 | myosin, heavy polypeptide 2, skeletal muscle, adult | G3UW82 | 5.18E-04 | -2.4 |
| 49 | S100b | S100 protein, beta polypeptide, neural | P50114 | 3.97E-03 | -2.4 |
| 50 | Cds2 | CDP-diacylglycerol synthase (phosphatidate cytidylyltransferase) 2 | Q99L43 | 1.01E-07 | -2.4 |

MGI gene symbols, protein names, UniProt accession numbers, Q-values, and fold-changes (FC) are shown. FC is the ratio (DTA/WT) of the averaged relative protein abundance in DRG. (-) indicates enrichment in $Na_V1.8^-$ nociceptors. Proteins were sorted by FC.

**Supplementary Table 3. Unmatched proteins (183) for which no corresponding transcripts could be found in the transcript list**

| No. | Gene Symbol | Description | Feature Type |
|---|---|---|---|
| 1 | Acad12 | acyl-Coenzyme A dehydrogenase family, member 12 | protein coding gene |
| 2 | Actbl2 | actin, beta-like 2 | protein coding gene |
| 3 | Adamts13 | a disintegrin-like and metallopeptidase (reprolysin type) with thrombospondin t | protein coding gene |
| 4 | AK157302 | cDNA sequence AK157302 | protein coding gene |
| 5 | Atp2b4 | ATPase, Ca++ transporting, plasma membrane 4 | protein coding gene |
| 6 | C7 | complement component 7 | protein coding gene |
| 7 | Ccdc190 | coiled-coil domain containing 190 | protein coding gene |
| 8 | Ccdc25 | coiled-coil domain containing 25 | protein coding gene |
| 9 | Cd63 | CD63 antigen | protein coding gene |
| 10 | Col28a1 | collagen, type XXVIII, alpha 1 | protein coding gene |
| 11 | Col6a6 | collagen, type VI, alpha 6 | protein coding gene |
| 12 | Cox7b | cytochrome c oxidase subunit 7B | protein coding gene |
| 13 | Cpt1b | carnitine palmitoyltransferase 1b, muscle | protein coding gene |
| 14 | D17H6S53E | DNA segment, Chr 17, human D6S53E | protein coding gene |
| 15 | Dnaaf3 | dynein, axonemal assembly factor 3 | protein coding gene |
| 16 | Ear10 | eosinophil-associated, ribonuclease A family, member 10 | protein coding gene |
| 17 | Ear2 | eosinophil-associated, ribonuclease A family, member 2 | protein coding gene |
| 18 | Ear6 | eosinophil-associated, ribonuclease A family, member 6 | protein coding gene |
| 19 | Eif1ad13 | eukaryotic translation initiation factor 1A domain containing 13 | protein coding gene |
| 20 | Eloc | elongin C | protein coding gene |
| 21 | Enpp7 | ectonucleotide pyrophosphatase/phosphodiesterase 7 | protein coding gene |
| 22 | Fam177a | family with sequence similarity 177, member A | protein coding gene |
| 23 | fh | fetal hematoma | heritable phenotypic marker |
| 24 | Frmpd4 | FERM and PDZ domain containing 4 | protein coding gene |
| 25 | Fv4 | Friend virus susceptibility 4 | heritable phenotypic marker |
| 26 | Gbp5 | guanylate binding protein 5 | protein coding gene |
| 27 | Gcat | glycine C-acetyltransferase (2-amino-3-ketobutyrate-coenzyme A ligase) | protein coding gene |
| 28 | Gm17669 | predicted gene, 17669 | pseudogene |
| 29 | Gm20425 | predicted gene 20425 | protein coding gene |
| 30 | Gm20431 | predicted gene 20431 | protein coding gene |

| | | | |
|---|---|---|---|
| 31 | Gm20521 | predicted gene 20521 | protein coding gene |
| 32 | Gm20547 | predicted gene 20547 | protein coding gene |
| 33 | Gm20671 | predicted gene 20671 | protein coding gene |
| 34 | Gm28040 | predicted gene, 28040 | protein coding gene |
| 35 | Gm43738 | predicted gene 43738 | protein coding gene |
| 36 | Gm49368 | predicted gene, 49368 | protein coding gene |
| 37 | Gm49486 | predicted gene, 49486 | protein coding gene |
| 38 | Gm7324 | predicted gene 7324 | protein coding gene |
| 39 | Gm9774 | predicted pseudogene 9774 | protein coding gene |
| 40 | Gng5 | guanine nucleotide binding protein (G protein), gamma 5 | protein coding gene |
| 41 | Gpx4 | glutathione peroxidase 4 | protein coding gene |
| 42 | Gvin1 | GTPase, very large interferon inducible 1 | protein coding gene |
| 43 | H1f1 | H1.1 linker histone, cluster member | protein coding gene |
| 44 | H1f3 | H1.3 linker histone, cluster member | protein coding gene |
| 45 | H1f5 | H1.5 linker histone, cluster member | protein coding gene |
| 46 | H2ac21 | H2A clustered histone 21 | protein coding gene |
| 47 | H2ac6 | H2A clustered histone 6 | protein coding gene |
| 48 | H2ax | H2A.X variant histone | protein coding gene |
| 49 | H2bc8 | H2B clustered histone 8 | protein coding gene |
| 50 | H2bu1-ps | H2B.U histone 1, pseudogene | pseudogene |
| 51 | H3f3c | H3 histone, family 3C | protein coding gene |
| 52 | Hba | hemoglobin alpha chain complex | complex/cluster/region |
| 53 | Hbb-bs | hemoglobin, beta adult s chain | protein coding gene |
| 54 | Hmgb1 | high mobility group box 1 | protein coding gene |
| 55 | Idi1 | isopentenyl-diphosphate delta isomerase | protein coding gene |
| 56 | Igha | immunoglobulin heavy constant alpha | gene segment |
| 57 | Ighg2b | immunoglobulin heavy constant gamma 2B | gene segment |
| 58 | Ighv1-18 | immunoglobulin heavy variable V1-18 | gene segment |
| 59 | Igkc | immunoglobulin kappa constant | gene segment |
| 60 | Igkv10-96 | immunoglobulin kappa variable 10-96 | gene segment |
| 61 | Igkv12-41 | immunoglobulin kappa chain variable 12-41 | gene segment |
| 62 | Igkv12-44 | immunoglobulin kappa variable 12-44 | gene segment |
| 63 | Igkv12-46 | immunoglobulin kappa variable 12-46 | gene segment |
| 64 | Igkv2-109 | immunoglobulin kappa variable 2-109 | gene segment |

| | | | |
|---|---|---|---|
| 65 | Igkv4-55 | immunoglobulin kappa variable 4-55 | gene segment |
| 66 | Igkv8-30 | immunoglobulin kappa chain variable 8-30 | gene segment |
| 67 | Impdh2 | inosine monophosphate dehydrogenase 2 | protein coding gene |
| 68 | Isg15 | ISG15 ubiquitin-like modifier | protein coding gene |
| 69 | Isoc2a | isochorismatase domain containing 2a | protein coding gene |
| 70 | Kirrel2 | kirre like nephrin family adhesion molecule 2 | protein coding gene |
| 71 | Klhl3 | kelch-like 3 | protein coding gene |
| 72 | Lrriq1 | leucine-rich repeats and IQ motif containing 1 | protein coding gene |
| 73 | Lsm5 | LSM5 homolog, U6 small nuclear RNA and mRNA degradation associated | protein coding gene |
| 74 | Ly6c2 | lymphocyte antigen 6 complex, locus C2 | protein coding gene |
| 75 | Mrps33 | mitochondrial ribosomal protein S33 | protein coding gene |
| 76 | mt-Atp6 | mitochondrially encoded ATP synthase 6 | protein coding gene |
| 77 | mt-Atp8 | mitochondrially encoded ATP synthase 8 | protein coding gene |
| 78 | mt-Co1 | mitochondrially encoded cytochrome c oxidase I | protein coding gene |
| 79 | mt-Co2 | mitochondrially encoded cytochrome c oxidase II | protein coding gene |
| 80 | mt-Co3 | mitochondrially encoded cytochrome c oxidase III | protein coding gene |
| 81 | mt-Nd1 | mitochondrially encoded NADH dehydrogenase 1 | protein coding gene |
| 82 | mt-Nd2 | mitochondrially encoded NADH dehydrogenase 2 | protein coding gene |
| 83 | mt-Nd4 | mitochondrially encoded NADH dehydrogenase 4 | protein coding gene |
| 84 | Mtrf1 | mitochondrial translational release factor 1 | protein coding gene |
| 85 | Mup2 | major urinary protein 2 | protein coding gene |
| 86 | Mup20 | major urinary protein 20 | protein coding gene |
| 87 | Mup8 | major urinary protein 8 | protein coding gene |
| 88 | Myl6b | myosin, light polypeptide 6B | protein coding gene |
| 89 | Naa12 | N(alpha)-acetyltransferase 12, NatA catalytic subunit | protein coding gene |
| 90 | Ndufa11 | NADH:ubiquinone oxidoreductase subunit A11 | protein coding gene |
| 91 | Ndufb1 | NADH:ubiquinone oxidoreductase subunit B1 | protein coding gene |
| 92 | Ndufb4 | NADH:ubiquinone oxidoreductase subunit B4 | protein coding gene |
| 93 | Ndufc2 | NADH:ubiquinone oxidoreductase subunit C2 | protein coding gene |
| 94 | Ndufs5 | NADH:ubiquinone oxidoreductase core subunit S5 | protein coding gene |
| 95 | Nme3 | NME/NM23 nucleoside diphosphate kinase 3 | protein coding gene |
| 96 | Npm1 | nucleophosmin 1 | protein coding gene |
| 97 | Nptxr | neuronal pentraxin receptor | protein coding gene |
| 98 | Ogdhl | oxoglutarate dehydrogenase-like | protein coding gene |

| | | | |
|---|---|---|---|
| 99 | Oscp1 | organic solute carrier partner 1 | protein coding gene |
| 100 | Pacsin2 | protein kinase C and casein kinase substrate in neurons 2 | protein coding gene |
| 101 | Palm3 | paralemmin 3 | protein coding gene |
| 102 | Pam16 | presequence translocase-asssociated motor 16 | protein coding gene |
| 103 | Pcdha4 | protocadherin alpha 4 | protein coding gene |
| 104 | Pdcd5 | programmed cell death 5 | protein coding gene |
| 105 | Phb | prohibitin | protein coding gene |
| 106 | Pin4 | peptidyl-prolyl cis/trans isomerase, NIMA-interacting, 4 (parvulin) | protein coding gene |
| 107 | Ppid | peptidylprolyl isomerase D (cyclophilin D) | protein coding gene |
| 108 | Ppp1cc | protein phosphatase 1 catalytic subunit gamma | protein coding gene |
| 109 | Prdx6 | peroxiredoxin 6 | protein coding gene |
| 110 | Ptges3 | prostaglandin E synthase 3 | protein coding gene |
| 111 | Qdpr | quinoid dihydropteridine reductase | protein coding gene |
| 112 | Ramac | RNA guanine-7 methyltransferase activating subunit | protein coding gene |
| 113 | Ran | RAN, member RAS oncogene family | protein coding gene |
| 114 | Rbm10 | RNA binding motif protein 10 | protein coding gene |
| 115 | Rbm4 | RNA binding motif protein 4 | protein coding gene |
| 116 | Rbx1 | ring-box 1 | protein coding gene |
| 117 | Retnlg | resistin like gamma | protein coding gene |
| 118 | Rnaset2b | ribonuclease T2B | protein coding gene |
| 119 | Rpl10 | ribosomal protein L10 | protein coding gene |
| 120 | Rpl17 | ribosomal protein L17 | protein coding gene |
| 121 | Rpl21 | ribosomal protein L21 | protein coding gene |
| 122 | Rpl22 | ribosomal protein L22 | protein coding gene |
| 123 | Rpl23 | ribosomal protein L23 | protein coding gene |
| 124 | Rpl23a | ribosomal protein L23A | protein coding gene |
| 125 | Rpl24 | ribosomal protein L24 | protein coding gene |
| 126 | Rpl28 | ribosomal protein L28 | protein coding gene |
| 127 | Rpl3 | ribosomal protein L3 | protein coding gene |
| 128 | Rpl30 | ribosomal protein L30 | protein coding gene |
| 129 | Rpl32 | ribosomal protein L32 | protein coding gene |
| 130 | Rpl34 | ribosomal protein L34 | protein coding gene |
| 131 | Rpl35a | ribosomal protein L35A | protein coding gene |
| 132 | Rpl7a | ribosomal protein L7A | protein coding gene |

| | | | |
|---|---|---|---|
| 133 | Rps13 | ribosomal protein S13 | protein coding gene |
| 134 | Rps14 | ribosomal protein S14 | protein coding gene |
| 135 | Rps18-ps5 | ribosomal protein S18, pseudogene 5 | pseudogene |
| 136 | Rps18-ps6 | ribosomal protein S18, pseudogene 6 | pseudogene |
| 137 | Rps26 | ribosomal protein S26 | protein coding gene |
| 138 | Rps27a | ribosomal protein S27A | protein coding gene |
| 139 | Rps3a2 | ribosomal protein S3A2 | pseudogene |
| 140 | Rpsa | ribosomal protein SA | protein coding gene |
| 141 | Rragb | Ras-related GTP binding B | protein coding gene |
| 142 | Samd4b | sterile alpha motif domain containing 4B | protein coding gene |
| 143 | Sap30bp | SAP30 binding protein | protein coding gene |
| 144 | Scamp3 | secretory carrier membrane protein 3 | protein coding gene |
| 145 | Scamp4 | secretory carrier membrane protein 4 | protein coding gene |
| 146 | Sec31b | Sec31 homolog B (S. cerevisiae) | protein coding gene |
| 147 | Sec61g | SEC61, gamma subunit | protein coding gene |
| 148 | Septin5 | septin 5 | protein coding gene |
| 149 | Serpina1d | serine (or cysteine) peptidase inhibitor, clade A, member 1D | protein coding gene |
| 150 | Serpina1e | serine (or cysteine) peptidase inhibitor, clade A, member 1E | protein coding gene |
| 151 | Sh3bp1 | SH3-domain binding protein 1 | protein coding gene |
| 152 | Slc10a5 | solute carrier family 10 (sodium/bile acid cotransporter family), member 5 | protein coding gene |
| 153 | Slc25a3 | solute carrier family 25 (mitochondrial carrier, phosphate carrier), member 3 | protein coding gene |
| 154 | Smim15 | small integral membrane protein 15 | protein coding gene |
| 155 | Sms | spermine synthase | protein coding gene |
| 156 | Snu13 | SNU13 homolog, small nuclear ribonucleoprotein (U4/U6.U5) | protein coding gene |
| 157 | Sra1 | steroid receptor RNA activator 1 | protein coding gene |
| 158 | Srp14 | signal recognition particle 14 | protein coding gene |
| 159 | Srsf10 | serine and arginine-rich splicing factor 10 | protein coding gene |
| 160 | Syt15 | synaptotagmin XV | protein coding gene |
| 161 | Taco1 | translational activator of mitochondrially encoded cytochrome c oxidase I | protein coding gene |
| 162 | Tbrg4 | transforming growth factor beta regulated gene 4 | protein coding gene |
| 163 | Tceal5 | transcription elongation factor A (SII)-like 5 | protein coding gene |
| 164 | Tcp11l1 | t-complex 11 like 1 | protein coding gene |
| 165 | Tm9sf1 | transmembrane 9 superfamily member 1 | protein coding gene |
| 166 | Tmem14c | transmembrane protein 14C | protein coding gene |

| | | | |
|---|---|---|---|
| 167 | Tmsb15l | thymosin beta 15b like | protein coding gene |
| 168 | Tomm5 | translocase of outer mitochondrial membrane 5 | protein coding gene |
| 169 | Tomm6 | translocase of outer mitochondrial membrane 6 | protein coding gene |
| 170 | Tradd | TNFRSF1A-associated via death domain | protein coding gene |
| 171 | Trappc10 | trafficking protein particle complex 10 | protein coding gene |
| 172 | Trex1 | three prime repair exonuclease 1 | protein coding gene |
| 173 | Trim72 | tripartite motif-containing 72 | protein coding gene |
| 174 | Tubal3 | tubulin, alpha-like 3 | protein coding gene |
| 175 | Twf1 | twinfilin actin binding protein 1 | protein coding gene |
| 176 | Ube2l3 | ubiquitin-conjugating enzyme E2L 3 | protein coding gene |
| 177 | Ubl4a | ubiquitin-like 4A | protein coding gene |
| 178 | Ubl5b | ubiquitin-like 5B | protein coding gene |
| 179 | Ugt1a6b | UDP glucuronosyltransferase 1 family, polypeptide A6B | protein coding gene |
| 180 | Uprt | uracil phosphoribosyltransferase | protein coding gene |
| 181 | Usp35 | ubiquitin specific peptidase 35 | protein coding gene |
| 182 | Vps28 | vacuolar protein sorting 28 | protein coding gene |
| 183 | Xkr7 | X-linked Kx blood group related 7 | protein coding gene |

#### Supplementary Table 4. $Na_V1.8^+$ nociceptor-enriched proteins (78) in DRG

| No. | Gene Symbol | Fold-Change (DTA/WT) | log2 (Fold-Change) | Q-value (DTA/WT) | - log10(Q-value) | Descriptions | UniProtIds | Feature Type |
|---|---|---|---|---|---|---|---|---|
| 1 | Acpp | -10.653613 | -3.413271 | 0.000000 | 6.494792 | Prostatic acid phosphatase;Isoform 2 of Pro | Q8CE08;Q8CE08 | protein coding gene |
| 2 | Actn3 | -2.212682 | -1.145796 | 0.004468 | 2.349896 | Alpha-actinin-3 | O88990 | protein coding gene |
| 3 | Agpat4 | -2.213337 | -1.146223 | 0.001414 | 2.849504 | 1-acyl-sn-glycerol-3-phosphate acyltransfer | Q8K4X7 | protein coding gene |
| 4 | Ankrd13d | -2.037347 | -1.026692 | 0.044554 | 1.351117 | Ankyrin repeat domain-containing protein 1 | Q6PD24 | protein coding gene |
| 5 | Arhgap28 | -4.594784 | -2.199997 | 0.004716 | 2.326391 | Rho GTPase-activating protein 28;Rho GTPa | E9Q642;Q8BN5 | protein coding gene |
| 6 | Atrnl1 | -2.764496 | -1.467016 | 0.006509 | 2.186507 | Attractin-like protein 1;Isoform 2 of Attract | Q6A051;Q6A05 | protein coding gene |
| 7 | AU040320 | -3.550775 | -1.828134 | 0.045500 | 1.341992 | Dyslexia-associated protein KIAA0319-like p | Q8K135;Q8K135 | protein coding gene |
| 8 | Avil | -2.045866 | -1.032712 | 0.000000 | 12.405248 | Advillin | O88398 | protein coding gene |
| 9 | Calca | -14.408184 | -3.848817 | 0.001364 | 2.865149 | Calcitonin gene-related peptide 1 | Q99JA0 | protein coding gene |
| 10 | Camk2a | -16.040381 | -4.003637 | 0.000000 | 16.261828 | Calcium/calmodulin-dependent protein kin | F8WIS9;P11798 | protein coding gene |
| 11 | Ccdc141 | -2.525474 | -1.336554 | 0.000243 | 3.614436 | Coiled-coil domain-containing 141 | A2AST1;E9Q8Q | protein coding gene |
| 12 | Cdc42 | -2.037211 | -1.026595 | 0.000173 | 3.762216 | Isoform 1 of Cell division control protein 42 | P60766-1 | protein coding gene |
| 13 | Cds2 | -2.367249 | -1.243211 | 0.000000 | 6.996414 | Phosphatidate cytidylyltransferase 2 | Q99L43 | protein coding gene |
| 14 | Chsy3 | -4.990650 | -2.319228 | 0.027187 | 1.565643 | Chondroitin sulfate synthase 3 | Q5DTK1 | protein coding gene |
| 15 | Clgn | -3.108210 | -1.636084 | 0.000000 | 7.650937 | Calmegin | P52194 | protein coding gene |
| 16 | Cpeb4 | -2.199585 | -1.137232 | 0.000088 | 4.055689 | Cytoplasmic polyadenylation element-bindi | Q5SU47;Q5SU4 | protein coding gene |
| 17 | Cpt1c | -2.501833 | -1.322986 | 0.013514 | 1.869219 | Carnitine O-palmitoyltransferase 1, brain is | Q8BGD5 | protein coding gene |
| 18 | Ctsl | -2.066212 | -1.046988 | 0.000175 | 3.757681 | Cathepsin L1 | P06797 | protein coding gene |
| 19 | )130043K22Ri | -3.225612 | -1.689573 | 0.008331 | 2.079321 | Dyslexia-associated protein KIAA0319 hom | Q5SZV5 | protein coding gene |
| 20 | Dgkg | -3.081118 | -1.623454 | 0.000018 | 4.750496 | Diacylglycerol kinase gamma | Q91WG7 | protein coding gene |
| 21 | Dgkh | -3.822114 | -1.934371 | 0.000000 | 23.087306 | Diacylglycerol kinase | A0A2I3BQ43;A0 | protein coding gene |
| 22 | Dgki | -5.568794 | -2.477365 | 0.000000 | 13.932921 | Diacylglycerol kinase | D3YWQ0;D3Z2V | protein coding gene |
| 23 | Dgkz | -3.696220 | -1.886050 | 0.000000 | 12.362249 | Diacylglycerol kinase;Diacylglycerol kinase;I | A2AHJ7;A2AHK | protein coding gene |
| 24 | Dhcr24 | -2.068076 | -1.048290 | 0.000007 | 5.136577 | Delta(24)-sterol reductase | Q8VCH6 | protein coding gene |
| 25 | Disp2 | -3.572671 | -1.837003 | 0.010579 | 1.975538 | Protein dispatched homolog 2 | Q8CIP5 | protein coding gene |
| 26 | Dok4 | -2.349947 | -1.232628 | 0.015896 | 1.798707 | Docking protein 4 | Q99KE3 | protein coding gene |
| 27 | Dolpp1 | -2.892900 | -1.532516 | 0.034682 | 1.459894 | Dolichyldiphosphatase 1 | Q9JMF7 | protein coding gene |
| 28 | Eml1 | -3.569070 | -1.835548 | 0.000000 | 22.780642 | Echinoderm microtubule-associated protei | Q05BC3;Q05BC | protein coding gene |
| 29 | Fam169a | -2.002464 | -1.001777 | 0.012126 | 1.916296 | Soluble lamin-associated protein of 75 kDa | Q5XG69 | protein coding gene |

| | | | | | | | | |
|---|---|---|---|---|---|---|---|---|
| 30 | Fxyd2 | -2.937812 | -1.554742 | 0.000000 | 6.673074 | Sodium/potassium-transporting ATPase sul | Q04646 | protein coding gene |
| 31 | Galnt17 | -3.005739 | -1.587720 | 0.000099 | 4.003520 | Polypeptide N-acetylgalactosaminyltransfe | Q7TT15 | protein coding gene |
| 32 | Gfra2 | -2.711791 | -1.439246 | 0.000002 | 5.712546 | GDNF family receptor alpha-2 | O08842 | protein coding gene |
| 33 | H3f3c | -2.200984 | -1.138148 | 0.017280 | 1.762466 | Histone H3.3C;Histone H3.3 | P02301;P84244 | protein coding gene |
| 34 | Hps1 | -2.413249 | -1.270977 | 0.031189 | 1.506003 | Hermansky-Pudlak syndrome 1 protein hon | A0A0R4J062;E9 | protein coding gene |
| 35 | Icmt | -2.168495 | -1.116694 | 0.021724 | 1.663059 | Protein-S-isoprenylcysteine O-methyltransf | Q3U4N2;Q9EQI | protein coding gene |
| 36 | Ids | -2.641196 | -1.401191 | 0.000012 | 4.915527 | Iduronate 2-sulfatase | Q08890 | protein coding gene |
| 37 | Kcnt1 | -3.865256 | -1.950564 | 0.000266 | 3.575844 | Potassium channel subfamily T member 1 | A0A0G2JER3;A0 | protein coding gene |
| 38 | Kctd16 | -2.686292 | -1.425616 | 0.004525 | 2.344349 | BTB/POZ domain-containing protein KCTD1 | Q5DTY9 | protein coding gene |
| 39 | Kif5a | -2.134441 | -1.093858 | 0.000000 | 9.859137 | Kinesin heavy chain isoform 5A | P33175 | protein coding gene |
| 40 | Klf1 | -2.333415 | -1.222443 | 0.000000 | 7.786259 | UPF0577 protein KIAA1324-like homolog;Is | Q3UZV7;Q3UZV | protein coding gene |
| 41 | Kndc1 | -7.688689 | -2.942738 | 0.014883 | 1.827323 | Kinase non-catalytic C-lobe domain-contain | Q0KK55 | protein coding gene |
| 42 | L1cam | -2.247797 | -1.168512 | 0.000000 | 10.705837 | Neural cell adhesion molecule L1;L1 cell adl | P11627;Q6PGJ3 | protein coding gene |
| 43 | Mbnl2 | -2.023854 | -1.017105 | 0.038673 | 1.412587 | Isoform 4 of Muscleblind-like protein 2 | Q8C181-4 | protein coding gene |
| 44 | Msmo1 | -2.016966 | -1.012187 | 0.026011 | 1.584842 | Methylsterol monooxygenase 1 | Q9CRA4 | protein coding gene |
| 45 | Myh1 | -2.181316 | -1.125199 | 0.000000 | 8.291537 | Myosin-1 | Q5SX40 | protein coding gene |
| 46 | Myh2 | -2.400951 | -1.263606 | 0.000518 | 3.285569 | MCG140437, isoform CRA_d | G3UW82 | protein coding gene |
| 47 | Myh4 | -2.637749 | -1.399307 | 0.003927 | 2.405896 | Myosin-4 | Q5SX39 | protein coding gene |
| 48 | Nedd4l | -4.087266 | -2.031136 | 0.000000 | 10.467408 | E3 ubiquitin-protein ligase NEDD4-like;E3 u | E9PXB7;Q8CFI0 | protein coding gene |
| 49 | Nipsnap3b | -2.288081 | -1.194138 | 0.002261 | 2.645630 | Protein NipSnap homolog 3B | Q9CQE1 | protein coding gene |
| 50 | Osbpl3 | -3.400133 | -1.765591 | 0.000000 | 24.647672 | Oxysterol-binding protein;Oxysterol-bindin | D3YTT6;Q9DBS | protein coding gene |
| 51 | P2rx3 | -6.823287 | -2.770467 | 0.000479 | 3.319696 | P2X purinoceptor;P2X purinoceptor;P2X pu | A2AW03;A2AW | protein coding gene |
| 52 | Pgap1 | -2.034158 | -1.024432 | 0.000000 | 6.853268 | GPI inositol-deacylase | Q3UUQ7 | protein coding gene |
| 53 | Phf24 | -4.387869 | -2.133521 | 0.000000 | 16.946360 | PHD finger protein 24 | Q80TL4 | protein coding gene |
| 54 | Pirt | -3.618590 | -1.855428 | 0.000003 | 5.601023 | Phosphoinositide-interacting protein | Q8BFY0 | protein coding gene |
| 55 | Plcb3 | -2.890539 | -1.531338 | 0.000000 | 25.464473 | 1-phosphatidylinositol 4,5-bisphosphate ph | P51432 | protein coding gene |
| 56 | Plcxd3 | -2.841155 | -1.506477 | 0.004504 | 2.346424 | MCG49978;PI-PLC X domain-containing pro | G3X9A7;Q8BLJ3 | protein coding gene |
| 57 | Pqlc3 | -2.247864 | -1.168555 | 0.000009 | 5.069180 | PQ-loop repeat-containing protein 3 | Q8C6U2 | protein coding gene |
| 58 | Prkcd | -3.215095 | -1.684861 | 0.000000 | 19.913865 | Protein kinase C delta type | P28867 | protein coding gene |
| 59 | Prkcq | -2.083968 | -1.059333 | 0.000023 | 4.637347 | Protein kinase C theta type | Q02111 | protein coding gene |
| 60 | Ptrh1 | -4.588019 | -2.197871 | 0.029506 | 1.530084 | Probable peptidyl-tRNA hydrolase | Q8BW00 | protein coding gene |
| 61 | Rgs10 | -2.926472 | -1.549163 | 0.000000 | 7.258250 | Regulator of G-protein signaling 10 | Q9CQE5 | protein coding gene |
| 62 | Rgs3 | -9.058443 | -3.179263 | 0.000000 | 7.342054 | Regulator of G-protein signaling 3;Isoform ! | Q9DC04;Q9DC0 | protein coding gene |

| | | | | | | | | |
|---|---|---|---|---|---|---|---|---|
| 63 | S100b | -2.367714 | -1.243495 | 0.003968 | 2.401436 | Protein S100-B | P50114 | protein coding gene |
| 64 | Scg3 | -3.268107 | -1.708455 | 0.000000 | 8.413934 | Secretogranin-3;Isoform 2 of Secretogranin | P47867;P47867 | protein coding gene |
| 65 | Scn10a | -4.983505 | -2.317161 | 0.000001 | 6.144607 | Sodium channel protein type 10 subunit alp | Q6QIY3;Q6QIY3 | protein coding gene |
| 66 | Scn11a | -123.965137 | -6.953791 | 0.000000 | 11.995004 | Sodium channel protein type 11 subunit alp | Q9R053 | protein coding gene |
| 67 | Scn7a | -2.091551 | -1.064573 | 0.000000 | 13.728999 | Sodium channel protein | B1AYL1 | protein coding gene |
| 68 | Slc36a1 | -4.689701 | -2.229496 | 0.035682 | 1.447556 | Proton-coupled amino acid transporter 1 | Q8K4D3 | protein coding gene |
| 69 | Sqle | -2.135400 | -1.094506 | 0.003543 | 2.450685 | Squalene monooxygenase | P52019 | protein coding gene |
| 70 | Srl | -2.900342 | -1.536223 | 0.029125 | 1.535734 | Sarcalumenin | Q7TQ48 | protein coding gene |
| 71 | St8sia3 | -2.976849 | -1.573786 | 0.006966 | 2.157034 | Sia-alpha-2,3-Gal-beta-1,4-GlcNAc-R:alpha | Q64689 | protein coding gene |
| 72 | Stoml1 | -2.073627 | -1.052157 | 0.034438 | 1.462959 | Stomatin-like protein 1 | A0A0B4J1F1;Q8 | protein coding gene |
| 73 | Tmem177 | -5.223718 | -2.385077 | 0.025942 | 1.585998 | Transmembrane protein 177 | Q8BPE4 | protein coding gene |
| 74 | Tmem87b | -2.030145 | -1.021583 | 0.048598 | 1.313380 | Transmembrane protein 87B;Isoform 2 of T | Q8BKU8;Q8BKU | protein coding gene |
| 75 | Tnni2 | -2.124342 | -1.087016 | 0.019079 | 1.719448 | Troponin I, fast skeletal muscle (Fragment); | A2A6K0;P13412 | protein coding gene |
| 76 | Tnnt3 | -2.361212 | -1.239528 | 0.022005 | 1.657482 | Troponin T, fast skeletal muscle;Troponin T | A0A0R4J1B1;A2 | protein coding gene |
| 77 | Trarg1 | -3.196428 | -1.676461 | 0.045627 | 1.340777 | Trafficking regulator of GLUT4 1 | Q8C838 | protein coding gene |
| 78 | Trpv1 | -4.578864 | -2.194990 | 0.003117 | 2.506291 | Transient receptor potential cation channel | Q704Y3;Q704Y3 | protein coding gene |

#### Supplementary Table 5. Proteins (47) enriched in $Na_V1.8^-$ cells of DRG

| No. | Gene Symbol | Fold-Change (DTA/WT) | log2 (Fold-Change) (DTA/WT) | Q-value (DTA/WT) | - log10(Q-value) (DTA/WT) | Description | UniProtIds | Feature Type |
|---|---|---|---|---|---|---|---|---|
| 1 | Adgra2 | 2.501760 | 1.322944 | 0.011314 | 1.946369 | adhesion G protein-coupled | J3QMG7;Q91ZV8 | protein coding gene |
| 2 | Ahsp | 2.527292 | 1.337593 | 0.000034 | 4.471755 | alpha hemoglobin stabilizing | Q9CY02 | protein coding gene |
| 3 | Ambp | 2.126609 | 1.088555 | 0.003678 | 2.434334 | alpha 1 microglobulin/bikun | Q07456 | protein coding gene |
| 4 | Cdh13 | 2.100334 | 1.070619 | 0.000078 | 4.108717 | cadherin 13 | Q9WTR5 | protein coding gene |
| 5 | Ces1d | 2.693832 | 1.429660 | 0.004366 | 2.359950 | carboxylesterase 1D | Q8VCT4 | protein coding gene |
| 6 | Cldn1 | 2.000652 | 1.000470 | 0.003758 | 2.425090 | claudin 1 | O88551 | protein coding gene |
| 7 | Cma1 | 2.235273 | 1.160451 | 0.010325 | 1.986098 | chymase 1, mast cell | A4QPC5;P21844 | protein coding gene |
| 8 | Col6a6 | 2.095371 | 1.067206 | 0.000000 | 18.481199 | collagen, type VI, alpha 6 | E9Q6A6;Q8C6K9 | protein coding gene |
| 9 | Cpa3 | 2.450266 | 1.292938 | 0.000000 | 25.600117 | carboxypeptidase A3, mast c | P15089 | protein coding gene |
| 10 | Csrp2 | 2.158743 | 1.110191 | 0.001298 | 2.886850 | cysteine and glycine-rich pro | P97314 | protein coding gene |
| 11 | Cxadr | 2.089279 | 1.063005 | 0.011552 | 1.937352 | coxsackie virus and adenovi | P97792 | protein coding gene |
| 12 | Cyp2e1 | 3.460293 | 1.790894 | 0.003896 | 2.409368 | cytochrome P450, family 2, s | Q05421 | protein coding gene |
| 13 | Cyp2f2 | 2.822907 | 1.497181 | 0.004185 | 2.378323 | cytochrome P450, family 2, s | P33267 | protein coding gene |
| 14 | Ddc | 3.080310 | 1.623075 | 0.002932 | 2.532840 | dopa decarboxylase | O88533 | protein coding gene |
| 15 | Eln | 2.101074 | 1.071127 | 0.032963 | 1.481979 | elastin | P54320 | protein coding gene |
| 16 | Emilin1 | 2.045258 | 1.032283 | 0.000682 | 3.166523 | elastin microfibril interfacer | Q99K41 | protein coding gene |
| 17 | Fga | 2.182949 | 1.126278 | 0.000000 | 8.484317 | fibrinogen alpha chain | E9PV24 | protein coding gene |
| 18 | Fgb | 2.049586 | 1.035332 | 0.000000 | 26.228837 | fibrinogen beta chain | Q8K0E8 | protein coding gene |
| 19 | Fhl2 | 2.066982 | 1.047526 | 0.000008 | 5.077995 | four and a half LIM domains | O70433 | protein coding gene |
| 20 | Hp | 5.604128 | 2.486490 | 0.000000 | 13.668275 | haptoglobin | Q61646 | protein coding gene |
| 21 | Hpx | 2.079276 | 1.056081 | 0.000000 | 16.946360 | hemopexin | Q91X72 | protein coding gene |
| 22 | Igf2bp3 | 2.035673 | 1.025506 | 0.005221 | 2.282216 | insulin-like growth factor 2 r | Q9CPN8 | protein coding gene |
| 23 | Ighg2b | 2.293875 | 1.197787 | 0.000000 | 11.101599 | immunoglobulin heavy cons | A0A075B5P3;A0A | gene segment |
| 24 | Ighm | 2.193225 | 1.133054 | 0.000000 | 7.257718 | immunoglobulin heavy cons | A0A075B5P6;A0A | gene segment |
| 25 | Ighv6-6 | 2.239117 | 1.162930 | 0.015587 | 1.807248 | immunoglobulin heavy varia | A0A075B5T3 | gene segment |
| 26 | Igkc | 2.109991 | 1.077237 | 0.000000 | 6.380062 | immunoglobulin kappa cons | P01837 | gene segment |
| 27 | Igkv8-30 | 3.937627 | 1.977326 | 0.007101 | 2.148703 | immunoglobulin kappa chai | A0A140T8M3 | gene segment |

| | | | | | | | | |
|---|---|---|---|---|---|---|---|---|
| 28 | Igsf5 | 2.346345 | 1.230415 | 0.000015 | 4.833359 | immunoglobulin superfamil | D3YXH0;P63054 | protein coding gene |
| 29 | Inpp5k | 2.172867 | 1.119600 | 0.033042 | 1.480940 | inositol polyphosphate 5-ph | Q8C5L6 | protein coding gene |
| 30 | Itih4 | 2.367900 | 1.243608 | 0.000000 | 21.231657 | inter alpha-trypsin inhibitor, | A6X935;E9PVD2, | protein coding gene |
| 31 | Kcnj13 | 2.280209 | 1.189166 | 0.000084 | 4.077021 | potassium inwardly-rectifyir | G3UX25;P86046 | protein coding gene |
| 32 | Lrg1 | 3.859911 | 1.948567 | 0.000000 | 16.271355 | leucine-rich alpha-2-glycopr | Q91XL1 | protein coding gene |
| 33 | Mcpt4 | 2.567234 | 1.360215 | 0.000060 | 4.224512 | mast cell protease 4 | P21812;Q3UN88 | protein coding gene |
| 34 | Mocs1 | 2.223527 | 1.152850 | 0.014919 | 1.826266 | molybdenum cofactor synth | Q5RKZ7 | protein coding gene |
| 35 | Morn4 | 4.749358 | 2.247732 | 0.024598 | 1.609094 | MORN repeat containing 4 | Q6PGF2 | protein coding gene |
| 36 | Myl6b | 2.757517 | 1.463370 | 0.026961 | 1.569257 | myosin, light polypeptide 6E | Q8CI43 | protein coding gene |
| 37 | Parp3 | 2.042327 | 1.030214 | 0.003211 | 2.493392 | poly (ADP-ribose) polymeras | Q3ULW8;Q3ULW | protein coding gene |
| 38 | Plin1 | 2.886457 | 1.529300 | 0.038724 | 1.412019 | perilipin 1 | Q8CGN5 | protein coding gene |
| 39 | Ptgds | 2.274722 | 1.185690 | 0.027920 | 1.554085 | prostaglandin D2 synthase ( | O09114 | protein coding gene |
| 40 | Saa1 | 5.198455 | 2.378083 | 0.018803 | 1.725766 | serum amyloid A 1 | P05366 | protein coding gene |
| 41 | Serpinb6b | 6.142915 | 2.618923 | 0.042171 | 1.374982 | serine (or cysteine) peptidas | O08804 | protein coding gene |
| 42 | Sik3 | 4.596257 | 2.200459 | 0.014443 | 1.840342 | SIK family kinase 3 | E9PU87;F6S7W6 | protein coding gene |
| 43 | Slc4a10 | 2.390808 | 1.257498 | 0.009725 | 2.012131 | solute carrier family 4, sodiu | B1AWV9;Q5DTL! | protein coding gene |
| 44 | Tgfbi | 2.046026 | 1.032824 | 0.000000 | 15.308543 | transforming growth factor, | P82198 | protein coding gene |
| 45 | Thbs1 | 2.167674 | 1.116148 | 0.000487 | 3.312828 | thrombospondin 1 | P35441;Q80YQ1 | protein coding gene |
| 46 | Thrsp | 2.107691 | 1.075663 | 0.006641 | 2.177798 | thyroid hormone responsive | Q62264 | protein coding gene |
| 47 | Tpsb2 | 2.452118 | 1.294028 | 0.000642 | 3.192645 | tryptase beta 2 | E9QJW9;P21845 | protein coding gene |

**Suppl. Table 6.** The expression (as indicated by MouseBrain.org) of representative candidates shared among Top-50 transcripts (Suppl. Table 1) and Top-50 proteins (Suppl. Table 2)

| Index | Name | Description | Scn10a | Scn11a | P2rx3 | Prkcd | Acpp |
|---|---|---|---|---|---|---|---|
| 198 | PSPEP8 | Peptidergic (TrpM8), DRG | 0.00 | 0.00 | 0.00 | 0.499 | 0.142 |
| 199 | PSPEP7 | Peptidergic (TrpM8), DRG | 0.0788 | 0.00 | 0.00 | 0.139 | 0.282 |
| 200 | PSPEP6 | Peptidergic (TrpM8), DRG | 0.00 | 0.0552 | 0.0368 | 0.333 | 0.0933 |
| 201 | PSPEP5 | Peptidergic (PEP1.2), DRG | 0.199 | 0.434 | 0.00 | 0.801 | 0.0324 |
| 202 | PSPEP2 | Peptidergic (PEP1.3), DRG | 1.56 | 0.852 | 0.0881 | 1.74 | 0.148 |
| 203 | PSPEP4 | Peptidergic (PEP1.1), DRG | 1.16 | 1.55 | 0.0595 | 1.71 | 0.0162 |
| 204 | PSPEP3 | Peptidergic (PEP1..4), DRG | 2.51 | 0.901 | 0.130 | 1.97 | 0.556 |
| 205 | PSPEP1 | Peptidergicv (PEP2), DRG | 3.28 | 0.513 | 0.141 | 1.55 | 0.116 |
| 206 | PSNF3 | Neurofilament (NF2/3), DRG | 0.613 | 0.290 | 0.290 | 0.0658 | 0.0973 |
| 207 | PSNF2 | Neurofilament (NF4/5), DRG | 0.0350 | 0.122 | 0.00 | 0.0696 | 0.0183 |
| 208 | PSNF1 | Neurofilament (NF1), DRG | 0.0263 | 0.0256 | 0.00 | 0.104 | 0.501 |
| 209 | PSNP1 | Non-peptidergic (TH), DRG | 1.08 | 0.667 | 0.0813 | 0.665 | 2.91 |
| 210 | PSNP2 | Non-peptidergic (NP1.1), DRG | 3.71 | 3.39 | 0.305 | 2.56 | 4.16 |
| 211 | PSNP3 | Non-peptidergic (NP1.2), DRG | 5.28 | 4.80 | 0.965 | 7.02 | 4.24 |
| 212 | PSNP4 | Non-peptidergic (NP2.1), DRG | 5.47 | 4.93 | 0.244 | 5.09 | 5.07 |
| 213 | PSNP5 | Non-peptidergic (NP2.2), DRG | 4.99 | 4.57 | 0.556 | 4.97 | 3.66 |
| 214 | PSNP6 | Non-peptidergic (NP3), DRG | 4.00 | 3.77 | 1.21 | 9.67 | 3.59 |
